# Pupil-DLC: an open-source deep learning pipeline for scalable, marker-less tracking of pupil dynamics across conscious and unconscious states

**DOI:** 10.64898/2026.01.18.700183

**Authors:** Parsa Seyfourian, Lydia C. Marks, Leslie D. Claar, Yasmeen Nahas, Miles Keating, Christof Koch, Irene Rembado

## Abstract

**Background:** Pupil diameter is a non-invasive biomarker of brain state, correlating with arousal, attention, cognitive processing, and consciousness. However, existing pupillometry software often lacks scalability and robustness across diverse experimental conditions and species.

**New method:** We introduce Pupil-DLC, an open-source, offline, DeepLabCut-based pipeline for scalable, marker-less pupil tracking, primarily designed for mice. Trained on 21,750 manually annotated frames from over 140 videos of head-fixed mice spanning wakefulness and drug-induced states, including psychedelics and anesthesia, the dataset was deliberately selected to maximize pupil size variability and model generalization. Pupil-DLC implements a dual-model architecture: a General Model (GM) for high-throughput analysis and an Individual Model (IM) for session-specific optimization.

**Results:** Pupil-DLC captures pupil dynamics across awake, psychedelic, and anesthetized conditions with high agreement with ground truth and equal tracking fidelity during active locomotion and quiet rest. Confidence metrics aligned with human frame quality assessments, enabling principled tuning of accuracy–retention trade-offs. As a secondary demonstration, Pupil-DLC extends to unseen human videos across diverse conditions and frame rates, including daylight and smartphone recordings, without retraining.

**Comparison with existing methods:** Pupil-DLC outperforms existing automated methods in accuracy and frame retention while maintaining computational efficiency comparable to real-time tools. These improvements stem from a learned keypoint-based representation robust to pupil shape variability, occlusions, reflections, and imaging artifacts. The GM/IM framework supports a tiered strategy balancing throughput and precision.

**Conclusions:** Pupil-DLC provides a reproducible, adaptable platform for quantifying pupil-linked brain state dynamics across experimental paradigms and species, bridging basic mouse neuroscience and translational human applications.

## Introduction

Changes in pupil diameter, when corrected for luminance and depth of focus, provide a robust index of internal brain state across species^1,2^. Pupil fluctuations track arousal, attention, emotional processing, cognitive load, and autonomic physiology, reflecting strong coupling between central and peripheral nervous system dynamics^3–9^. During monotonous tasks, such as simulated driving, pupil diameter shows large fluctuations that accompany drowsiness and performance lapses, whereas emotionally salient states including fear, stress, or anxiety elicit pronounced pupillary responses^8,10,11^. Even in the absence of overt behavioral or emotional changes, pupil dynamics track substates of wakefulness^7,12^ and capture moment-to-moment variations in cognitive effort, decision timing, and perceptual selection^13^, and can reveal which of multiple competing sensory streams is being attended^14–16^. Clinically, pupillometry is increasingly used as a non-invasive biomarker of impairment or loss of consciousness, enabling objective neurological monitoring and prognostic assessment^17–19^. Together, these findings establish pupil dynamics as a powerful, non-invasive biomarker of neural state.

Despite the potential role of pupil dynamics as a biomarker for different experimental and clinical conditions, this measurement is often limited by proprietary or rigid software pipelines that lack scalability, reproducibility, and flexibility across experimental paradigms. Many existing approaches are sensitive to lighting variability, require manual correction, or fail to integrate with modern deep learning frameworks^20^. Recent deep learning-based tools, such as MEYE^21^, have made important progress toward open-source, cross-species pupillometry; however, they lack a framework for diverse-state training and session-specific model optimization. To address these limitations, we present a customizable, open-source pupil-tracking pipeline built on DeepLabCut^22^ and trained on 21,750 manually annotated frames collected from over 140 videos of head-fixed and behaving mice spanning diverse drug-induced states of consciousness, including psychedelics and anesthesia. These were deliberately selected to maximize pupil size variability in the training dataset, thereby improving model generalization across conditions. Our method enables high-resolution, marker-less pupil tracking with post-processing tools that improve accuracy and reduce noise; it also generalizes robustly to videos from another species not included in the annotated dataset, such as humans. While Pupil-DLC is primarily designed for mouse neuroscience experiments, this mouse-to-human generalization serves as a secondary demonstration of the pipeline’s robustness to diverse recording conditions. By combining this computational method with the sensitivity of pupillometry to brain state, this platform provides a flexible foundation for studying arousal, cognition, conscious and unconscious states, and neural dynamics in mice and humans as an offline, high-throughput analysis tool.

## Methods

Experimental procedures followed closely to those described previously in Claar, Rembado, et al., 2023 and Russo et al., 2025^23,24^. The methods are summarized below, and additional details are provided for novel procedures.

### Mice

C57BL/6J wild-type (n=69) were purchased from *The Jackson Laboratory* (JAX stock #000664) on postnatal day 28-35. Mice were housed in the Allen Institute animal facility and used in accordance with protocols approved by the Allen Institute’s Institutional Animal Care and Use Committee (IACUC) under protocol 25105, 2212 and 2003. All mice were housed in a shared facility with room temperatures between 20 and 22°C and humidity between 30 and 70%. Mice were maintained on a reverse 12-h light cycle, and were single-housed following surgery, with *ad libitum* access to food and water.

### Headframe Implantation Surgery

Prior to undergoing experimental procedures, all mice were implanted with a titanium headframe to facilitate head fixation *in vivo* ^25^. Surgery took place when mice were 4-11 weeks old. Pre-operative injections of dexamethasone (3-4 mg/kg, IM) and ceftriaxone (100-125 mg/kg, SC) were administered one to three hours prior to surgery. Mice were deeply anesthetized with isoflurane (5%) in an induction chamber and then placed on a stereotaxic rig (Model# 1900, KOPF; Tujunga, CA) and maintained at a surgical level of anesthesia using isoflurane (1.5-2.5%) via nose cone. Breathing was monitored and body temperature was maintained at 37.5 °C with a heating pad under the animal (TC-1000 temperature controller, CWE, Inc.). Ocular lubricant (Systane, Alcon Inc., Geneva, Switzerland) was applied to the eyes during anesthesia to maintain hydration. During surgery, skin was removed to expose the skull, and the skull was leveled with reference to bregma. Using white C&B Metabond (Parkell, Inc., Edgewood, New York), a custom titanium headframe was secured to the skull, and the rest of the exposed skull was covered. After surgery, the mouse was given an injection of lactated Ringer’s solution (up to 1mL, SC) and was placed on a heating pad to recover. All animals received analgesics and antibiotics for two days post-surgery.

### Habituation

Five days after headframe implantation surgery, all mice began a 3-week training schedule. Researchers first habituated mice to handling for 2 days. On subsequent days, mice were placed on a running wheel where the headframe was fixed in place with two sets of screws^25^. During training, mice were kept head-fixed on the running wheel for an increasing amount of time, starting with 5-minute sessions and ending with 90-minute sessions. At the end of the second week of training, the mice scheduled to receive psilocybin on the experimental day began receiving 2 intraperitoneal (IP) injections of saline during each training session (method described below). All training and experimental sessions were performed during the dark cycle.

### Experimental Sessions

After mice completed 3 weeks of habituation to the experimental setup, they underwent one recording a day for three consecutive days. Half of the animals received isoflurane on the first day and psilocybin on the second day, and the other half experienced the reverse order. The urethane recording always took place on the third day as it was considered a terminal procedure under our IACUC protocol. During each recording session, mice were head-fixed on a wheel, and data was collected for 45-120 minutes. On the isoflurane and psilocybin recording days, there was a period of 30-60 minutes of baseline activity recorded during the awake state before administering the drug, followed by 30-60 minutes after drug administration. On the urethane day, there was no awake state baseline period pre-sedation given that the urethane was administered intravenously via a catheter inserted in its tail vein before setting the animal on the rig (see below for details). On the experimental day the angular position of the running wheel was acquired by a dedicated computer with a National Instruments card acquiring digital inputs at 100 kHz, which was considered the master clock. A 32-bit digital “barcode” was sent with an Arduino Uno (DEV-11021, SparkFun Electronics, Niwot, Colorado) every 30 seconds to synchronize all devices^26^.

### Psilocybin and Ketanserin Administration

Psilocybin was administered via an intraperitoneal (IP) injection while the animal was head-fixed on the wheel. The researcher scuffed the loose skin over the mouse’s back and immobilized the two hind legs to prevent the mouse from moving. While the animal was restrained, the researcher administered the injection into the peritoneal cavity on the lower right quadrant of the mouse’s abdomen. During psilocybin recordings, mice received one injection of sterile saline (1 mg/kg, IP) followed 10-12 minutes later by an injection of psilocybin (1 mg/kg, IP; Usona Institute). During psilocybin+ketanserin recordings, the saline injection was substituted with a ketanserin (1 mg/kg, IP; S006, Sigma-Aldrich) injection.

### Anesthesia Scale

To ensure a reliable and reproducible state of consciousness during both isoflurane and urethane states, a behavioral scale was developed based on Devor and Zalkind, 2001 to evaluate depth of anesthesia^27^. The scale quantifies the mouse’s posture and response to noxious stimuli on a scale of 0-4, where a higher number corresponded to a minimal or no response, generally observed during deep levels of anesthesia. The four tests included an assessment of muscle tone and voluntary movement (posture), withdrawal reflexes of tail and foot when a firm pinch was applied (300-500 grams of force) through a rodent pincher analgesia meter (IITC Life Science), and response to the presence of a noxious alcohol swab held near the nose. The sum of the scores from each individual test ranged from 0-16 and represented 4 states on the continuum of consciousness. On the assumption that a loss of responsiveness corresponds to a loss of consciousness, a summed score < 6 corresponded to a “conscious” state, while a score of 6-10 represented a “mildly anesthetized” state. To be considered unconscious, a mouse needed a score ≥ 11, which corresponds to either a “completely unconscious” state (11-13) or a “deep anesthesia” state (14-16).

### Isoflurane Administration

To induce isoflurane anesthesia while head-fixed on the wheel, the mouse received inhalant isoflurane delivered through a small tube placed in front of the mouse’s nose. After 2-5 minutes at 5% isoflurane, the flow was reduced to 1-2% for the remainder of the recording. 10 minutes after the start of isoflurane, the mouse was given a score based on the anesthesia scale. If the score was lower than 11, the percentage of isoflurane was increased, and the test was administered again 10 minutes later until the mouse scored 11 or higher on the scale. Throughout the recording, the precise concentration of isoflurane being administered was recorded. During anesthesia, a heating pad was placed under the animal and set at 37.5 °C to maintain thermoregulation. At the end of the recording, the isoflurane was switched off, and the mouse recovered before being placed back in its home cage.

### Urethane Administration

Prior to intravenous (IV) injection of urethane, the mouse was warmed under an infrared heat lamp for 4 minutes to increase body temperature and promote vasodilation. The animal was then anesthetized using isoflurane inhalant (3-5%) and placed on a gel heating pad to maintain thermoregulation. The tail was dipped in warm water to increase vasodilation, and an alcohol swab was administered to sterilize the tail and increase visibility of the lateral tail vein. A beveled 31G needle attached to infusion catheter tubing (SAI, Infusion Technologies) was held parallel to the mouse’s tail and inserted into the lateral vein. Correct placement of the needle was confirmed by the presence of blood in the infusion tube and smooth injection of fluid without any resistance on the plunger of the syringe. Once the needle was in the vein, *Vetbond Tissue Adhesive* (Patterson Veterinary, Loveland, CO) was applied to the exposed area of the needle to hold it in place. Isoflurane was discontinued, and the mouse was moved to a restrainer where the urethane (1.5-1.8 mg/kg) was administered via the tail vein catheter at a rate of 0.02 mL/min. The depth of anesthesia was tested 60-90 minutes after the initial urethane injection and if the animal scored 10 or below, supplemental doses (20% of initial dose) were administered (up to 60% of initial dose) until the animal achieved a score of 11 or higher. When the IV injection was unsuccessful (10 out of 32 animals), urethane (1.5-1.8 mg/kg) was administered in two IP injections, each containing half of the full dose, 15 minutes apart. During anesthesia, a heating pad was placed under the animal and set at 37.5 °C to maintain thermoregulation.

### Pupil Video Recordings

Videos of the right eye were acquired with an infrared camera (Mako G-032B PoE 552; Proximity Series 130mm/0.73x; MAX: 000F315C1496) at a sampling rate of 30Hz with a resolution of 640×480 pixels. The camera was pointing to an infrared dichroic mirror positioned at around 9 cm from the camera, reflecting the right eye of the animal positioned 3 cm away from the mirror (Fig. 1A). The pupil radius ranged from a minimum of 10 pixels (radius) to a maximum of 125 pixels, with 77 pixels corresponding to 1 mm for our specific setup. A black curtain was lowered over the front of the rig, placing the mouse in complete darkness.

**Figure 1.**
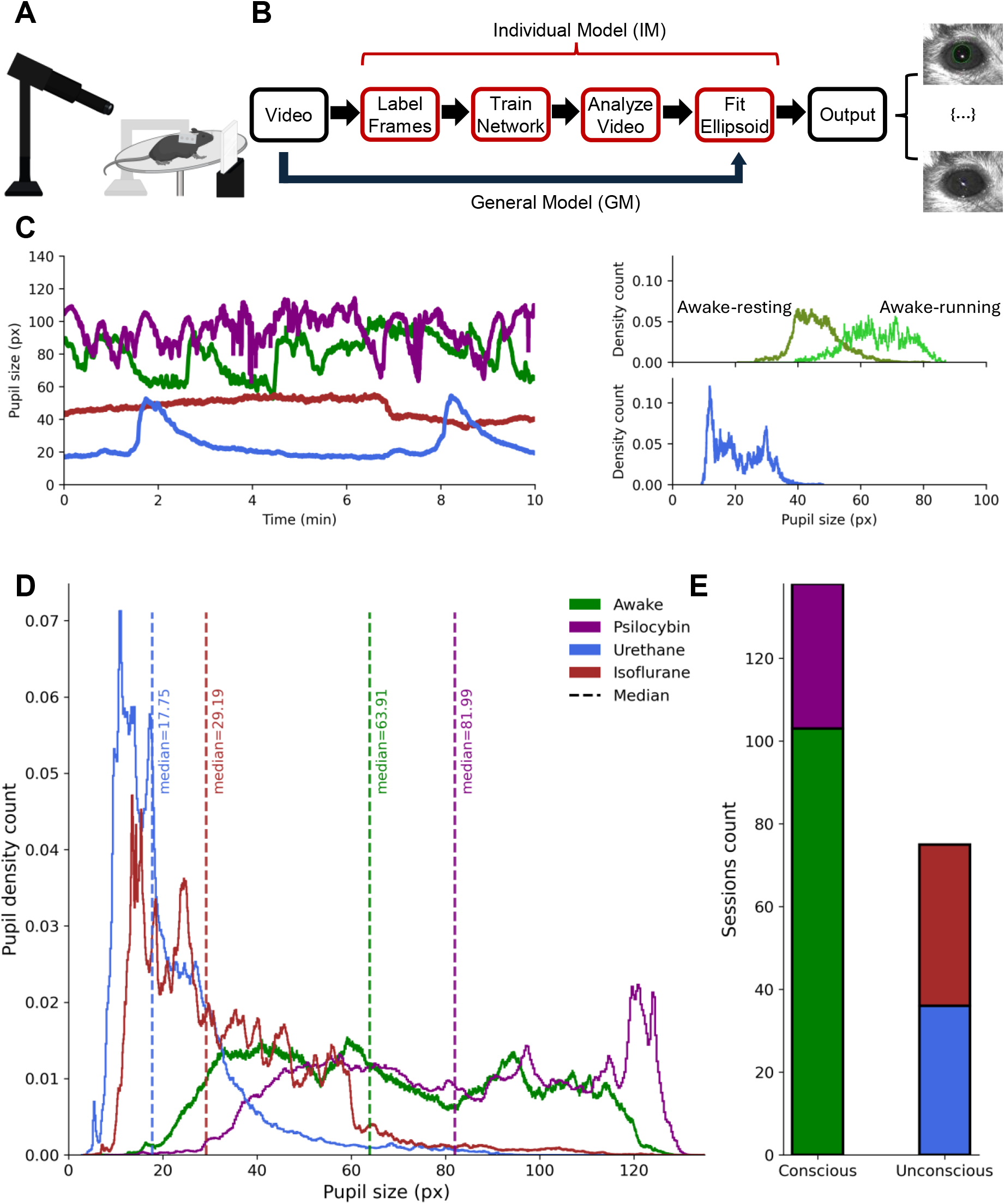
Pupillometry measurements in head-fixed mice undergoing different conscious and unconscious states. A) Experimental setup schematic of the head-fixed mouse on a rotating wheel (to monitor locomotion). The infrared camera points to a dichroic mirror reflecting the right eye of the animal utilized to track pupillometry. B) Pupil-DLC pipeline overview. Two main paths are provided by the pipeline: a general model (GM), trained on 21,750 manually annotated frames from over 140 videos of 69 mice, and an individual model (IM) trained on 150 frames of any one session. C) Left: Time courses of pupil size (in pixels, px) over a 10-minute recording session under four conditions: awake (green), after administration of psilocybin (purple), under urethane anesthesia (blue), and under isoflurane anesthesia (brown). Right: Probability density distributions of pupillometry across awake-resting, awake-running, and urethane states from a representative mouse. D) Probability density distributions of pupillometry across conscious and unconscious states in 51 mice. E) Summary bar-plot showing the total number of sessions in each state used to train the Pupil-DLC general model pooled over 140 videos from 69 mice. Note that for our mouse setup, the minimal pupil diameter of 10 pixels corresponds to 1/8 of a mm.

### Pupil Video Processing

We employed *DeepLabCut* (DLC, https://www.mackenziemathislab.org/deeplabcut), a non-commercial marker-less pose estimation toolbox based on deep convolutional neural networks^22^, to track pupil dynamics of mouse eye videos. DLC leverages transfer learning on deep neural networks (ResNet-50 backbone) to track user-defined keypoints with near human accuracy. To train the network, each frame was annotated with 12 equidistant points around the pupil’s contour, arranged in a clockwise orientation, and 4 additional reference points along the horizontal and vertical axes of the eye, identifying the medial and lateral commissures of each frame. These annotations served as training data for two model types: a General Model (**GM**) and an Individual-specific Model (**IM**). Pupil-DLC is designed as an offline analysis pipeline; all video processing is performed post-acquisition.

### DLC General Model Training

The GM of pupil-DLC was trained on a large, diverse dataset comprising 21,750 manually annotated frames selected from over 140 videos of different head-fixed mice under distinct conscious and unconscious states, including: urethane anesthesia, isoflurane anesthesia, awake, psilocybin, and psilocybin with ketanserin pretreatment. This training dataset included both active (e.g., running) and quiescent (e.g., resting) conscious states, thereby equipping the model with the ability to generalize across different eye sizes and shapes. GM training was performed using DLC’s ResNet-50 architecture for 1,000,000 iterations. The GM allows one to analyze new pupil videos without requiring per-session retraining, offering fast, high-throughput and reproducible pupil tracking across experimental paradigms.

### DLC Individual Model Training

For experiments requiring animal or session-specific precision, a tailored IM was used. For each new video, 150 frames were randomly sampled throughout the whole recording and manually annotated using the same 12-point pupil scheme described above. A separate ResNet-50 model was then fine-tuned for 500,000 iterations. This approach ensures high accuracy even in videos with atypical lighting, eye positioning, or artifact presence. The resulting model was then used to analyze the full video.

### DLC Fine-tuning Integration

To address cases where the General Model does not fully generalize to a new recording session, such as atypical lighting conditions, camera positioning, or inter-individual differences in eye anatomy, we implemented a session-specific fine-tuning mode (GM-FT) as an alternative to the GM and IM workflows. Rather than initializing a ResNet-50 network from ImageNet-pretrained weights, GM-FT starts directly from the fully trained GM checkpoint (1,000,000 iterations) and continues training on a small set of session-specific frames. For each new session, frames were manually selected and annotated using the same labeling protocol described above. Fine-tuning ran for up to 200,000 additional iterations under a three-stage learning rate schedule designed to preserve the GM’s learned representations while allowing adaptation to session-specific appearance statistics: 5×10^-4^ for iterations 1–10,000; 10^-4^ for iterations 10,001–100,000; and 5×10^-5^ thereafter. These rates are approximately 10-fold lower than those used during GM training, reducing the risk of catastrophic forgetting. Because GM-FT operates on the GM weights rather than the full training corpus, no labeled data from prior sessions is required and retraining time is substantially reduced relative to the IM. In our testing, we terminated the training after 50,000 iterations.

### Post-training and Ellipse Fitting

Once the full video was processed through the IM or GM, DeepLabCut produced, for each frame, the predicted pupil boundary keypoints (*x*_*i*_, *y*_*i*_) together with a confidence value *p*_*i*_. Only keypoints with confidence *p*_*i*_ ≥ 0.9 were retained for geometric estimation. For each frame, ellipse fitting proceeded only when at least six high confidence pupil boundary points were available. Frames that did not meet these criteria were excluded from ellipse fitting and all ellipse derived outputs for those frames were recorded as ‘Not a Number’ (NaN).

The pupil contour in each frame was approximated by fitting a single ellipse to the retained boundary points using a least squares estimator. An ellipse in the image plane is fully defined by five geometric parameters: the center coordinates (*x*_*c*_, *y*_*c*_), the semi major and semi minor radii (*a, b*) and the orientation angle *θ*. Because five independent parameters must be estimated, a minimum of five non-collinear points is required to uniquely determine an ellipse. In practice, using six or more points yields a reliable approach that improves robustness to noise and local boundary deviations.

Ellipse fitting was performed using the *EllipseModel* implementation in scikit-image^28^, which estimates ellipse parameters by minimizing the algebraic distance between the observed boundary points and the implicit ellipse model. Given a set of *n* ≥ 5 observed points (*x*_*i*_, *y*_*i*_), the ellipse is represented as a conic section

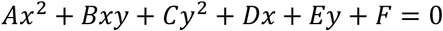

subject to the constraint that the conic corresponds to an ellipse. The model parameters are estimated using a direct least squares solution, after which the conic representation is converted into the geometric ellipse parameters (*x*_*c*_, *y*_*c*_, *a, b, θ*).

From the fitted ellipse, pupil size was quantified using the ellipse radii. In all primary analyses we reported pupil size as the pupil radius defined as the largest of the two semi axis lengths:

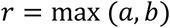

which provides a robust estimate of pupil size, insensitive to small deviations from circularity. Pupil diameter is approximated as 2*r* when required, and pupil area as *A* = π(*r*)^2.

For training the models and fitting the ellipse, a Windows 11 Pro desktop computer with the following specifics was used: AMD Ryzen Threadripper 3960X 24-Core CPU @ 3.80 GHz; RAM: 128GB; Graphics card: 2x NVIDIA GeForce RTX 3090 Ti.

### Inference Speed Optimization

To maximize throughput during offline video analysis, DLC’s default serial inference pipeline was replaced with a custom threaded implementation incorporating three complementary optimizations. First, a dedicated producer thread decodes and preprocesses video frames concurrently with GPU inference, eliminating the CPU–GPU serialization inherent in DLC’s default inference function and maximizing GPU utilization. Decoded frames are buffered into a bounded queue of up to eight pre-loaded batches, ensuring the GPU is never stalled awaiting input. Second, TensorFlow’s XLA (Accelerated Linear Algebra) JIT compiler was enabled at the highest optimization level (global_jit_level = ON_2) by injecting a modified ConfigProto into DLC’s GPU session setup. XLA fuses ResNet-50’s approximately 150 individual CUDA kernel launches into fewer, more efficient operations, substantially reducing kernel dispatch overhead. Automatic mixed precision (auto_mixed_precision = ON) was additionally enabled, allowing eligible ResNet-50 convolutions to execute in FP16 on the Ampere tensor cores of the RTX 3090 Ti. Third, an optional ONNX Runtime backend was implemented: when a trained DLC model has been converted to ONNX format, inference is routed through ONNX Runtime’s CUDA Execution Provider, which applies its own operator fusion and bypasses TensorFlow’s session overhead entirely; XLA serves as the fallback when no ONNX model is available. Across all inference paths, a vectorized NumPy heatmap postprocessing step replaces DLC’s per-joint Python loops with a single batched argmax and location-refinement operation. Together, these optimizations achieve inference throughput of up to 350 Hz when processing frames in batches of 64 on the hardware described above, enabling rapid analysis of multi-hour recordings without modification to DLC model weights or training procedure.

### Accuracy Assessment and Comparison with Existing Tools

To assess the accuracy of the outputs from either GM or IM, we compared the extracted pupil diameters with ground-truth (GT) manually measured pupil diameters obtained using ImageJ^29^. Notably, due to the manual nature of ground-truth assessment, these measurements are inherently subject to variability on the order of a few pixels. Moreover, we compared the outputs of our pipeline with the output of EyeLoop^30^ (EL), an open-source, Python-based, real-time eye-tracking toolkit designed for high-speed pupil detection using a vectorized contour-detection algorithm, that operates entirely on consumer-grade CPUs. Its modular architecture allows integration of custom importers and extractors, supports multiple species (including rodents), and is optimized for closed- and open-loop experimental paradigms. MEYE^21^ is an open-source, deep learning-based web application for pupillometry, compatible with infrared webcams and validated in both mice and humans. OpenIris^31,32^ (OI) is an open-source, infrared-based eye-tracking system that detects the pupil boundary via edge detection and tracks gaze using the double Purkinje image method, validated in non-human primates at sampling rates up to 500 Hz. In our study, all three tools served as benchmarks to evaluate Pupil-DLC performance in terms of accuracy, frame retention, and processing speed.

Statistical differences between estimates were assessed using a Student’s *t*-test when assumptions of normality and homogeneity of variance were satisfied, as verified by Shapiro–Wilk and Levene’s tests, respectively. When the normality assumption was not met, differences were evaluated using the Mann– Whitney U test.

## Results

### Pupil-DLC training dataset spans the full range of mouse pupil dynamics across conscious and unconscious states

We trained Pupil-DLC on a large and diverse library of pupil videos from head-fixed mice (Figure 1A) spanning multiple drug-induced states of consciousness, deliberately selected to maximize pupil size variability in the training set. Across conditions, pupil size (reported as the largest radius of the estimated ellipses, see Methods) exhibited the expected state-dependent differences, consistent with established literature. During awake and psilocybin conditions, pupils were larger on average and showed pronounced temporal fluctuations throughout the recordings, with a more dilated pupil during running compared to resting conditions (Figure 1C). In contrast, urethane and isoflurane anesthesia were associated with smaller pupil diameters and markedly reduced variability (Figure 1C). During urethane sedation, pupil size alternated between two distinct substates: a more aroused state characterized by pupil dilation and a less aroused state marked by sustained constriction (Figure 1C, D), consistent with previous reports^33^. These patterns were evident at the level of individual animals (Figure 1C) and remained robust when pooled across sessions and subjects (Figure 1D).

Probability density distributions of pupil diameter further confirmed these well-established effects, with conscious states exhibiting broader distributions shifted toward larger diameters, whereas unconscious states showed narrower distributions concentrated around smaller pupil sizes (Figure 1D). These distributions were consistent across mice, indicating that the observed effects were not driven by individual variability. Together, these results demonstrate that the Pupil-DLC pipeline reliably captures physiologically meaningful pupil dynamics across distinct levels of consciousness and underscores the breadth of the dataset used to train GM. In total, the dataset comprised 95 recordings during awake condition before drug administration, 35 recordings collected during isoflurane anesthesia, 32 during urethane sedation, and 35 following psilocybin administration (with or without ketanserin pre-treatment) (Figure 1E).

### Pupil-DLC achieves high accuracy relative to human ground truth and existing methods

To assess measurement accuracy, pupil size estimates obtained with the Pupil-DLC GM and IM were compared against human-annotated ground truth and three automated methods: EyeLoop, MEYE, and OpenIris. Across extended recordings, both Pupil-DLC models closely tracked GT measurements over time, faithfully capturing rapid pupil dynamics and remaining aligned across behavioral transitions. When comparing the raw pupil traces detected by the different methods, EyeLoop, MEYE, and OpenIris produced noisier traces with systematic deviations from GT (Figure 2A). Quantitative analyses confirmed these observations: median pupil size estimates from both Pupil-DLC models were substantially closer to GT values than those obtained with any of the three benchmark methods, with error distributions showing minimal deviation from GT for Pupil-DLC and markedly larger discrepancies for the alternatives (mean±STD difference in pixels to GT for GM: 1.28±1.09; IM: 1.22±1.04; EL: 23.81±16.85; OI: 27.43±23.11; MEYE: 35.96±19.73; Figure 2B). Notably, while EyeLoop, OpenIris, GM and IM detected the pupil in nearly all frames, MEYE showed substantially higher frame loss, retaining only approximately 50% of frames (Figure 2C).

**Figure 2.**
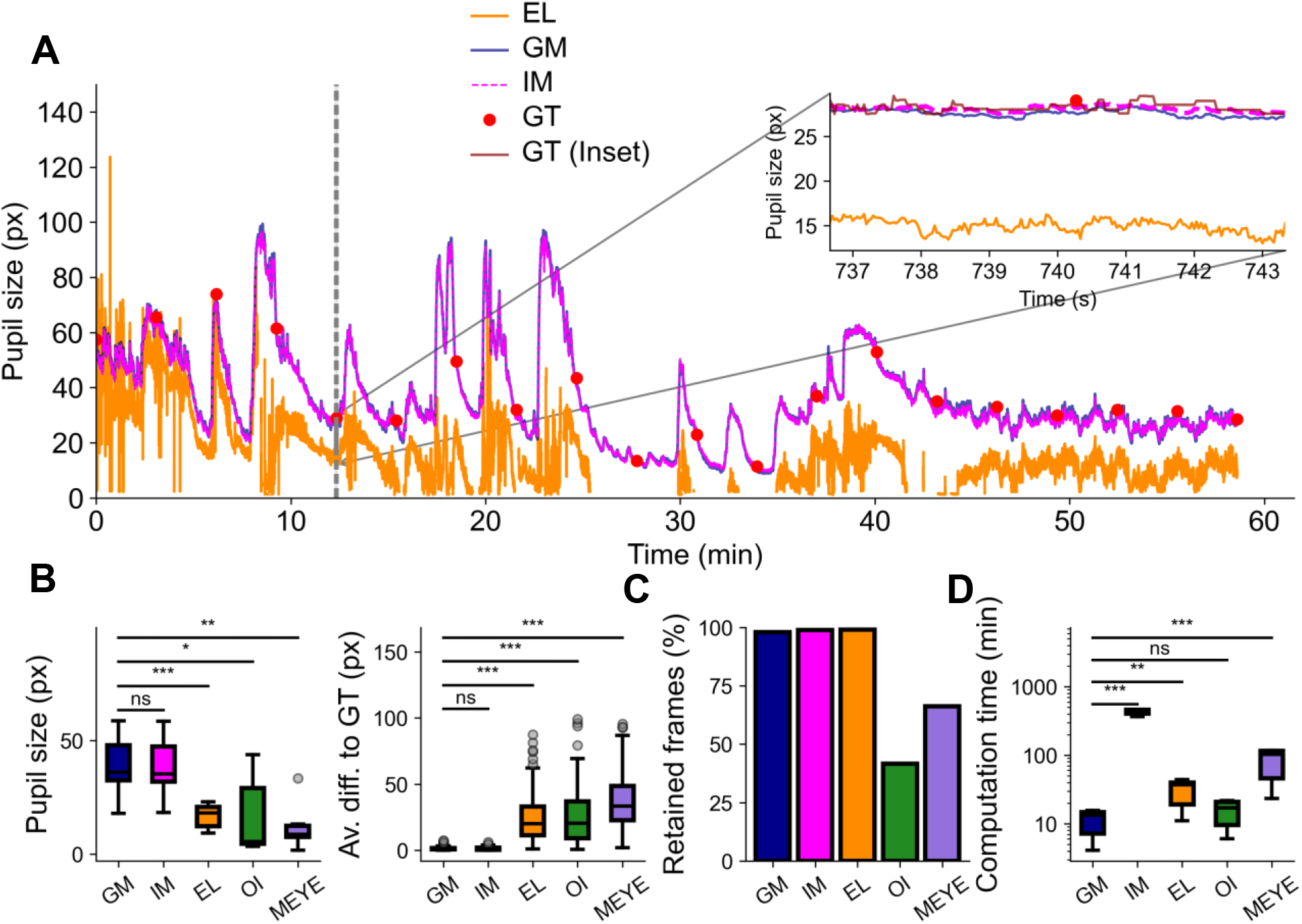
Performance evaluation of Pupil-DLC pipeline in measuring pupillometry across states compared to other approaches. A) Representative 60-minute pupillometry trace obtained from EyeLoop (EL, orange), General Model (GM, blue), and Individual Model (IM, magenta), compared to human-measured ground truth (GT, red). The inset shows a zoomed-in segment (6.67 seconds corresponding to 200 frames) to illustrate the close agreement between GM and IM with GT. B) Left: Absolute pupil size values (pixels) measured by GM, IM, EL, OpenIris (OI), and MEYE. Statistical test: Student t-test and Mann-Whitney U test. Right: Difference between GT measurements of frames compared to other methods. Statistical test: Mann-Whitney U test. C) Percentage of retained frames (i.e., frames in which the algorithm measured the pupil) for each method. D) Computation time per video for each method; y-axis is log-scale. Statistical test: Mann-Whitney U test. Statistical significance: n.s. p>0.05; * p<0.05; ** p<0.01; *** p<0.001.

IM’s superior accuracy incurred a ∼100-fold increase in computational cost. In contrast, GM preserved computational efficiency comparable to OpenIris (OI) while maintaining high accuracy (mean±STD minutes for GM: 11.40±4.16; IM: 423.84±33.53; EL: 31.00±11.50; OI: 15.52±5.85; MEYE: 84.58±36.39; Figure 2D). Together, these results demonstrate that Pupil-DLC enables accurate, robust, and computationally efficient pupil tracking over long-duration recordings.

### Pupil-DLC confidence metrics reliably capture measurement quality and accuracy

We next examined how the internal confidence estimates provided by the Pupil-DLC pipeline relate to measurement quality and accuracy. Across all frames, both GM and IM assigned consistently high confidence values, indicating reliable pupil detection (mean±STD confidence value for GM: 0.97±0.035; IM: 0.98±0.033; Figure 3A). Model confidence values closely matched human-based assessments of frame quality. Frames labeled as high quality by human observers were associated with higher confidence scores, whereas lower-quality frames showed systematically reduced confidence, validating the confidence metric as a meaningful proxy for visual measurement reliability (Figure 3B). Moreover, GM confidence values were not significantly different between resting and running conditions (Mann-Whitney U test, p>0.05 for both; Figure 3C), demonstrating that Pupil-DLC tracks pupil dynamics with equal fidelity during active locomotion and quiet rest.

**Figure 3.**
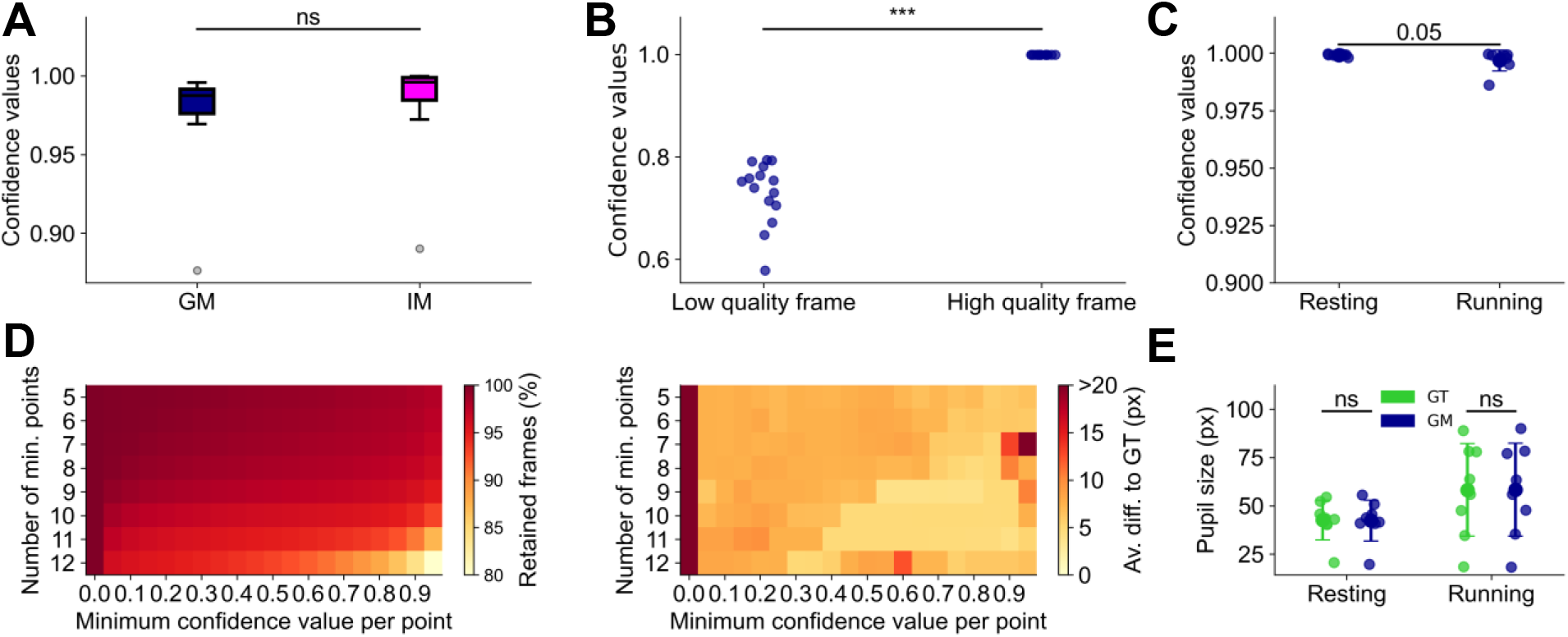
Performance evaluation as function of Pupil-DLC pipeline quality parameters. A) Confidence values assigned by the General Model (GM, blue) and the Individual Model (IM, magenta) across all frames averaged per video (n = 9 videos). B) GM confidence values as function of binary human-based frame quality assessment (n = 15 frames per quality category). C) GM confidence values as a function of resting and running frames averaged per video (n = 8 videos). D) Left: Percentage of frames retained based on the minimum number of annotated points as a function of confidence value per point. Right: Pixel difference to GT as a function of the ellipse fitting parameters (number of minimum points and minimum confidence value per point). Statistical test: Mann-Whitney U test. E) Absolute pupil size values (pixels) as a function of resting or running frames averaged per video, comparing against ground truth (GT). Statistical significance: n.s. p>0.05; * p<0.05; ** p<0.01; *** p<0.001.

Systematic variation of ellipse-fitting confidence thresholds revealed a clear trade-off between frame retention and measurement accuracy. By adjusting restraining parameters as defined by the minimum number of points required for ellipse fitting and the confidence threshold assigned to each point, we evaluated performance in terms of frame loss and deviation from ground truth. Increasing confidence thresholds and the minimum number of points reduced the number of retained frames (i.e. increased frame loss) but resulted in smaller average pixel deviation from ground truth, whereas more permissive thresholds increased frame retention at the expense of greater measurement error (Figure 3D). To further validate pipeline robustness across behavioral states, we compared tracking performance during resting and running conditions against GT and found that Pupil-DLC maintained accurate tracking in both (Figure 3E). Together, these results demonstrate that Pupil-DLC enables flexible tuning of analysis parameters to balance data completeness and accuracy according to experimental needs and confirm the robustness of Pupil-DLC across the full range of behavioral states relevant to the pupil–arousal literature.

### Pupil-DLC generalizes to unseen human pupil videos across recording conditions

Finally, we evaluated whether Pupil-DLC pipeline generalizes beyond its training domain by applying it to human pupil recordings acquired under infrared illumination and publicly available^34^ (Figure 4A). The dataset was recorded using a state-of-the-art dark-pupil, head-mounted eye tracker equipped with a *PointGrey Chameleon3* USB3.0 camera, modified to capture high-speed video at 95 frames per second. Recordings were collected across a variety of everyday indoor and outdoor environments, encompassing diverse illumination conditions (natural and artificial lighting) and a wide range of camera–eye angles. Some participants wore contact lenses or glasses, introducing realistic challenges such as reflections and partial occlusions, thereby increasing both the diversity and ecological validity of the dataset. We randomly selected ten human pupil recordings of various quality from this dataset to test Pupil-DLC performance. Importantly, this evaluation is entirely zero-shot: no human data were used at any stage of model training or validation. When video quality was high and reflections and occlusions were minimal, GM reliably detected and tracked the pupil in these human videos despite substantial variability in eye morphology and recording conditions, producing stable ellipse fits across frames (Figure 4A). Under these good conditions, both GM and IM yielded pupil size estimates that closely matched ground truth values (mean±STD pupil size value in pixels for GM: 57.55±11.15; IM: 57.50±11.09; GT: 59.18±11.64; Figure 4A). In contrast, for very low-quality videos at the limit of human visibility, GM failed to detect the pupil entirely (Figure 4B). However, IM still provided estimates closely aligned with ground truth, with a mean difference of only 3 pixels, well within the variability of the ground truth assessment (mean±STD difference from GT in pixels for IM: 3.35±2.81; GM-FT: 3.42±2.82; Figure 4B). Fine-tuning GM on a small (n = 150) set of session-specific frames fully recovered tracking performance, with GM-FT achieving accuracy comparable to IM (Figure 4B). To further assess pipeline generalizability, we evaluated pupil tracking on two static human color videos recorded under natural lighting using smartphones (Figure 4C). The videos differed in duration (10 vs. 30 seconds) and sampling rate (60 vs. 30 Hz). As expected, GM failed to track the pupil under these conditions, whereas IM closely matched GT. Fine-tuning GM on 150 session-specific frames per video allowed GM-FT to outperform GM and achieve accuracy comparable to IM (mean±STD deviation from GT, px: GM, 11.32±7.39; IM, 0.89±0.79; GM-FT, 0.71±0.75; Figure 4C). We next applied the same approach to a color video recorded at 24 Hz during head and camera motion (Figure 4D). GM again showed substantially higher tracking error; both IM and GM-FT tracked the pupil accurately, though GM-FT error was moderately higher than IM (mean±STD deviation from GT, px: GM, 4.92±5.10; IM, 0.62±0.47; GM-FT, 1.22±0.64; Figure 4D).

**Figure 4.**
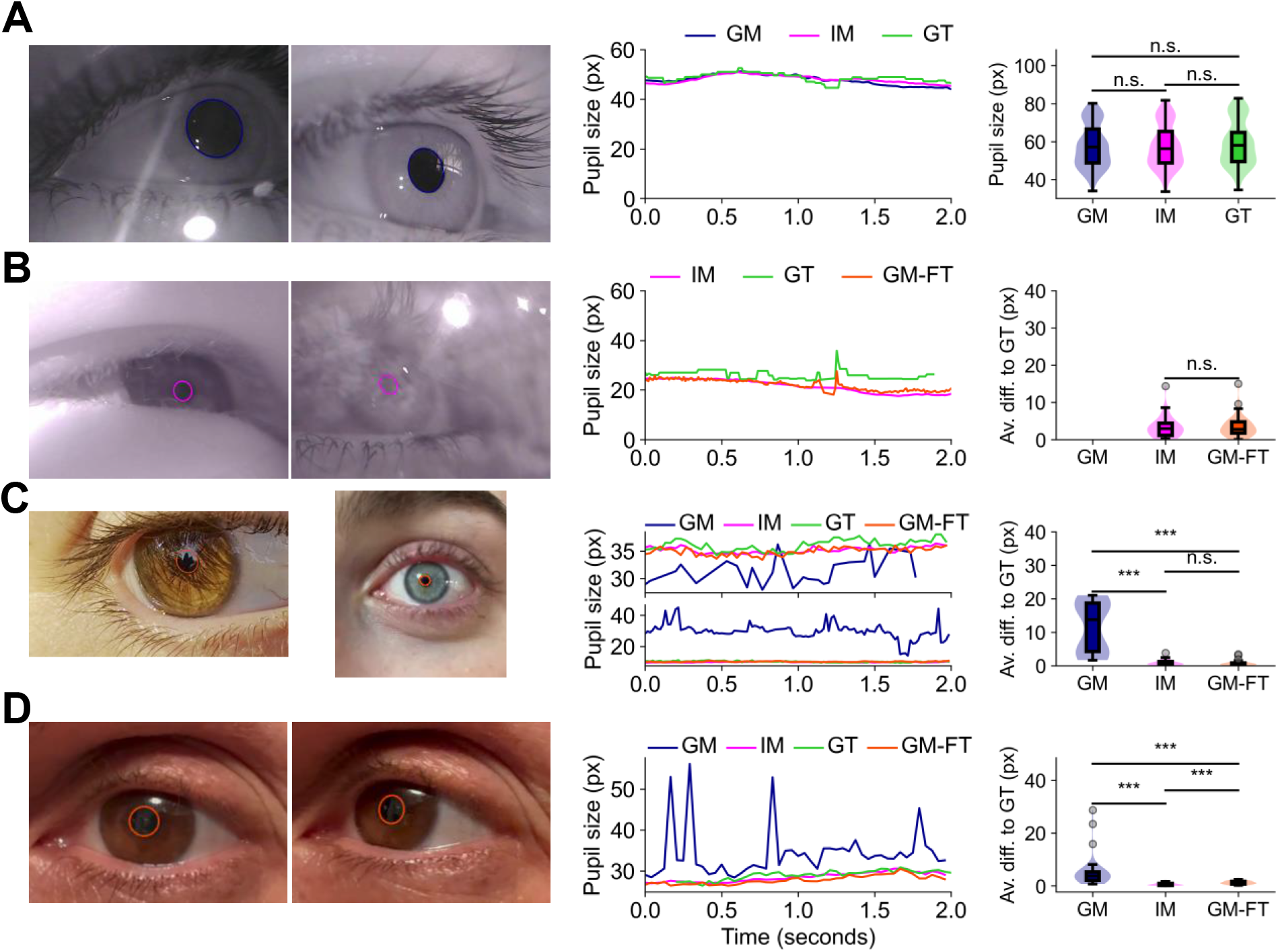
Pupil-DLC generalizes to unseen human pupil videos. A) Left: Representative frames from two good quality human pupil video recordings^34^, showing the pupil tracked by Pupil-DLC (GM) (dark blue ellipse overlaid). Middle: Short time-series (2 seconds, 180 frames) of estimated pupil size (in pixels) by Pupil-DLC (GM, dark blue; IM, magenta), compared to ground truth (GT, green). Right: Pupil size estimation by GM, IM and GT across eight good quality human pupil videos (n = 20 frames per video). B) Left: Representative frames from two bad quality human pupil video recordings^34^, showing the pupil tracked by Pupil-DLC (IM) (magenta ellipse overlaid). Middle: Short time-series (2 seconds, 180 frames) of estimated pupil size (in pixels) by Pupil-DLC (IM, magenta), and fine-tuned GM (GM-FT, orange), compared to ground truth (GT, green). Right: Pupil size estimation difference from GT by IM, and GM-FT across 2 bad quality human pupil videos (n = 20 frames per video). Note GM failed to track the pupil. C) Left: Representative frames from two static self-recorded human pupil videos, showing the pupil tracked by Pupil-DLC fine-tuned GM (GM-FT) (orange ellipse overlaid). Middle: Short time-series (2 seconds, 60 and 120 frames) of estimated pupil size (in pixels) by Pupil-DLC (GM, dark blue; IM, magenta; GM-FT, orange; GT, green). Right: Pupil size estimation difference from GT by GM, IM, and GM-FT across 2 human pupil videos (n = 20 frames per video). Statistical test: Mann-Whitney U test. D) Left: Representative frames from a self-recorded human pupil video captured during head and camera motion, with the pupil contour detected by GM-FT overlaid (orange ellipse). Middle: Two-second time series (48 frames) of pupil size estimates (px) from GM (dark blue), IM (magenta), GM-FT (orange), and GT (green). Right: Average deviation from GT for GM, IM, and GM-FT across one human pupil video (n = 20 GT-annotated frames). Statistical test: Mann– Whitney U test. n.s. p > 0.05; * p < 0.05; ** p < 0.01; *** p < 0.001.

Together, these results demonstrate that GM trained exclusively on mouse data generalizes to previously unseen recordings from human subjects without additional retraining when video quality closely matches the original training conditions. Under lower-quality or heterogeneous recording conditions, IM provides a reliable alternative, and fine-tuned GM with a small, video-specific dataset substantially extends its tracking fidelity. Collectively, these findings demonstrate that, as a secondary generalizability test, Pupil-DLC extends robustly to human recordings across diverse conditions, supporting its potential applicability beyond the mouse neuroscience context for which it was primarily designed.

## Discussion

In this study, we introduce *Pupil-DLC*, an open-source, DeepLabCut-based pipeline designed to overcome key limitations of existing pupillometry software and enable accurate extraction of pupil dynamics from video recordings. Pupil-DLC is trained on a large dataset of head-fixed mouse pupil videos spanning multiple drug-induced states of consciousness, with the explicit goal of providing scalable, marker-less tracking that generalizes across experimental conditions and video quality.

As expected, Pupil-DLC robustly captures state-dependent pupil dynamics across awake, psychedelic, and anesthetized conditions, consistent with the established view that pupil diameter reflects brain state, autonomic regulation, and state of consciousness^3–7,9^. In our dataset, pupil size distributions differed systematically between conscious and unconscious states, where more aroused states, such as running, are associated with a more dilated pupil compared to resting states, and urethane sedation exhibited alternating substates of dilation and constriction (Figure 1), in line with prior reports^12,33^.

Beyond capturing biologically meaningful structure, Pupil-DLC achieved high agreement with human-annotated ground truth. The present work builds on prior deep-learning-based pupillometry tools, most notably MEYE^21^, which introduced open-source, infrared-compatible pupil tracking across mice and humans. While MEYE represented an important step toward accessible cross-species pupillometry, Pupil-DLC offers several distinct contributions: a DeepLabCut-based keypoint representation that explicitly models the pupil boundary as geometric landmarks, providing robustness to partial occlusions, corneal reflections, and shape deformations; a GM/IM dual-model framework balancing high-throughput processing against session-specific precision; and training on a uniquely diverse dataset spanning multiple drug-induced states of consciousness, maximizing generalization across the full range of pupil sizes encountered in systems neuroscience. These design choices are reflected in the quantitative comparison: Pupil-DLC outperformed EyeLoop, OpenIris^31,32^, and MEYE in both accuracy and frame retention (Figure 2). It is worth noting that EyeLoop was designed primarily for real-time, closed-loop control and is therefore included here as a speed benchmark rather than a direct accuracy competitor. Across heterogeneous conditions, EyeLoop, OpenIris, and MEYE produced noisier estimates with systematic deviations from ground truth, whereas both GM and IM maintained high accuracy (Figure 2B). Importantly, Pupil-DLC is currently designed as an offline pipeline; nevertheless, the computational efficiency of the GM, comparable to EyeLoop, suggests online tracking is a feasible future extension that we are actively exploring.

A key practical contribution of this framework is the explicit separation between a GM, optimized for high-throughput analysis, and an IM, designed for session-specific optimization. Note that IM can be sufficiently trained on only 150 frames, in videos that can last over an hour or more, depending on the quality variability of the recording. GM is trained to analyze new videos without per-session retraining, enabling rapid, scalable, and reproducible processing across experiments (Figure 1B). However, in cases where recording conditions or video quality deviate substantially from the training data, performance of the GM may degrade. To address this, Pupil-DLC provides an IM that can be fine-tuned to new recordings, achieving high accuracy at the cost of increased computational time (Figure 2D). In practice, this design supports a tiered analysis strategy in which GM is used for large-scale screening and routine processing, while IM is reserved for challenging datasets or analyses requiring maximal precision.

An additional strength of Pupil-DLC is the availability of interpretable confidence metrics that closely align with human judgments of frame quality and can be leveraged to tune analysis stringency (Figure 3). Confidence values corresponded well with binary human-based assessments of frame quality, and systematic variation of ellipse-fitting thresholds revealed a clear trade-off between measurement accuracy and frame retention (Figure 3). This provides a principled framework for parameter selection depending on whether the priority is maximizing data yield or maximizing agreement with ground truth. Such flexibility is particularly valuable for long-duration recordings, where missing data can bias brain-state estimates, and for studies requiring stringent filtering to resolve fine-grained pupil dynamics. Furthermore, GM confidence values and tracking accuracy were statistically indistinguishable between resting and running epochs (Figure 3), confirming that the pipeline maintains equal measurement fidelity across behaviorally distinct states.

An unexpected and notable result was the cross-species generalization observed when applying the Pupil-DLC pipeline, trained exclusively on mouse data, to human pupil recordings acquired under infrared illumination and publicly available^34^. Importantly, this generalization is entirely zero-shot: no human data were used at any stage of model training or validation prior to this evaluation. Despite substantial differences in eye morphology and recording conditions, GM produced stable ellipse fits and pupil size estimates that closely matched human-annotated ground truth across multiple high-quality human videos (Figure 4A). In contrast, for poor-quality videos in which pupil detection was challenged by reflections and occlusions, GM failed to detect the pupil. However, IM successfully tracked the pupil even under these extreme conditions, deviating from manually estimated ground truth (itself subject to variability) by an average of only 3 pixels (Figure 4B). We further validated Pupil-DLC on three self-recorded human color videos spanning both stationary and dynamic (head-motion) recording conditions under natural lighting. While the GM failed, IM closely matched ground truth, and fine-tuned GM (GM-FT) on a small subset of frames restored performance to levels comparable to IM (Figure 4). These results highlight the flexibility of Pupil-DLC in bridging controlled laboratory recordings and real-world human data. The ability of a model trained exclusively on mouse pupil videos to generalize to human recordings underscores shared visual and geometric features across species when imaging conditions are comparable. Importantly, the complementary roles of GM and IM, together with rapid retraining using minimal data (GM-FT), provide a practical strategy for maintaining high tracking fidelity under heterogeneous recording conditions. Furthermore, Pupil-DLC infers pupil size on a frame-by-frame basis, making the pipeline entirely independent of video frame rate. This is demonstrated by its successful application to human recordings acquired at 95 frames per second and to self-recorded videos at 24, 30 and 60 frames per second, confirming compatibility with a broad range of camera frame rates beyond the 30 Hz used in our mouse recordings. This adaptability positions Pupil-DLC as a scalable tool for extending pupillometry beyond tightly controlled settings and into diverse experimental and translational contexts.

A few limitations should be acknowledged. First, the present study focuses on head-fixed recordings acquired under specific infrared imaging geometry. Performance may differ in freely moving conditions, at extreme gaze angles, or under variable illumination. However, the availability of the IM provides a flexible mechanism to adapt the pipeline to new recording conditions through targeted retraining. Second, pupil size outputs are reported in pixel units. While videos from any setup can be directly analyzed, conversion to physical units and comparisons across rigs require calibration of camera geometry and appropriate scaling. Physical unit conversion requires knowledge of the camera magnification, the distance between camera and eye, and the pixel size of the sensor. As a reference, in our specific setup 77 pixels correspond to 1 mm. We recommend that users perform a one-time calibration using an object of known physical size placed at the same distance as the eye.

In summary, Pupil-DLC is an open-source, scalable pupillometry pipeline primarily designed for mouse neuroscience that combines high accuracy, minimal frame loss, and tunable quality control, while also demonstrating robust generalizability to human eye videos as a secondary finding. By lowering technical barriers to robust pupil tracking, this framework enables reproducible measurement of pupil-linked brain state dynamics across diverse experimental contexts, spanning basic neuroscience in mice and translational applications in humans.

## Code and Data Availability

Pupil-DLC is freely available online: https://github.com/Valyriverse/Pupil-DLC.

A selection of representative unprocessed mouse pupil videos and the manually annotated frames used to train the General Model (GM) of *Pupil-DLC* are available in the Figshare repository: Manually annotated frames: Seyfourian P., et al. (2026), Pupil-DLC: an open-source deep learning pipeline for scalable, marker-less tracking of pupil dynamics across conscious and unconscious states (https://doi.org/10.6084/m9.figshare.31282714).

## Acknowledgments

We sincerely thank Peter Ledochowitsch for generous feedback and extended discussions. We thank the Allen Institute Animal Care, the Neurosurgery and Behavior, and the Lab Animal Services teams for mouse husbandry, care, and habituation; the Allen Institute Manufacturing and Process Engineering team for experimental hardware and software support. We wish to thank the Allen Institute founder, Paul G. Allen, for his vision, encouragement, and support.

We gratefully acknowledge funding from the Tiny Blue Dot Foundation (Santa Monica, California), the Templeton World Charity Foundation (TWCF-2022-30262; I.R.), and the Allen Institute.

